# Structural insights into the mechanism of nucleotide regulation of pancreatic K_ATP_ channel

**DOI:** 10.1101/2021.11.29.470334

**Authors:** Mengmeng Wang, Jing-Xiang Wu, Dian Ding, Xinli Duan, Songling Ma, Lipeng Lai, Lei Chen

## Abstract

ATP-sensitive potassium channels (K_ATP_) are metabolic sensors that convert the intracellular ATP/ADP ratio to the excitability of cells. They are involved in many physiological processes and implicated in several human diseases. Here we present the cryo-EM structures of the pancreatic K_ATP_ channel in both the closed state and the pre-open state, resolved in the same sample. The nucleotides bind at the inhibitory sites of the Kir6.2 channel in the closed state but not in the pre-open state. Structural comparisons reveal the mechanism for ATP inhibition and Mg-ADP activation, two fundamental properties of K_ATP_ channels. Moreover, the structure also uncovers the activation mechanism of diazoxide-type K_ATP_ openers.

## Introduction

The activity of K_ATP_ channels is inhibited by cytosolic ATP and activated by Mg-ADP^1^. The opening of K_ATP_ channels leads to the hyperpolarization of the cell, while the inhibition of K_ATP_ results in depolarization^2^. Therefore, K_ATP_ channels translate the cellular metabolic status into the excitability of the plasma membrane to control the electrical activity of the cell^1^. Because of its unique properties, K_ATP_ channels play essential roles in many key physiological processes, such as hormone secretion^2^, cardiac precondition^3^, and vasodilation ^4^. The genetic mutations of genes encoding K_ATP_ channels lead to a spectrum of diseases, ranging from metabolic syndrome to cardiovascular diseases and CNS disorders, including neonatal diabetes^5^, hyperinsulinaemic hypoglycemia of infancy ^5^, dilated cardiomyopathy ^6^, familial atrial fibrillation ^7^, Cantu syndrome ^8, 9^ and intellectual disability myopathy syndrome ^10^. K_ATP_ channels are also important drug targets. K_ATP_ inhibitors promote insulin release for the treatment of diabetes. These drugs include glibenclamide (GBM) and repaglinide (RPG), the so-called insulin secretagogues ^11^. K_ATP_ activators (K_ATP_ openers) are used to pharmacologically activate K_ATP_ channels in clinic ^12^. Diazoxide is an oral K_ATP_ opener and has been used in the treatment of hypoglycemia and hypertension for nearly half a century^12^.

Functional K_ATP_ channels are hetero-octamers composed of four Kir6 and four SUR subunits^13^. Kir6 are inward rectifier potassium channels that require PI(4,5)P_2_ for maximum activity^14–18^. It harbors the nucleotide-binding pocket which can bind the inhibitory ATP, and also ADP to a lesser extent ^14, 19^. SUR subunits are ABC transporter-like proteins that undergo Mg-nucleotide-dependent conformational changes^20^. The SUR subunits bind activating Mg- ADP and drugs, including insulin secretagogues and K_ATP_ openers ^11, 21^. Recent advances in cryo-EM structure determination of K_ATP_ channels in the presence of different ligand combinations by three groups have provided instrumental information about how K_ATP_ channels are assembled from individual subunits, how inhibitory ATP binds the channel, and how chemically distinct insulin secretagogues bind at SUR subunits^22–30^. Moreover, the conformational changes of the SUR1 subunit upon Mg-nucleotide binding have been visualized^22, 26^. Despite the progress, the fundamental questions about how K_ATP_ channels work, including the mechanism of ATP inhibition and Mg-ADP activation, remain elusive due to the lack of structures of K_ATP_ in the open state. To answer these outstanding questions, we sought to obtain the K_ATP_ channel in the open state.

## Results

### Structure determination

PI(4,5)P_2_ is important for K_ATP_ channel activity ^14–18^. However, previous trials of supplementing soluble PI(4,5)P_2_ analog PI(4,5)P_2_diC_8_ into K_ATP_ cryo-EM sample failed to stabilize the channel in the open state and no PI(4,5)P_2_diC_8_ density was observed^22, 26^. In contrast, there are several structures of other Kir family members in complex with PI(4,5)P_2_diC_8_ available, including Kir2.2^31^ or Kir3.2^32^. Therefore, we hypothesized the affinity of PI(4,5)P_2_diC_8_ for Kir6.2 might be lower than those of Kir2.2 or Kir3.2. In agreement with is, sequence alignments showed several positively charged residues at the PI(4,5)P_2_ binding pocket of Kir2.2 or Kir3.2 are replaced by non-charged polar residues in Kir6 channels (Fig. S1a). Particularly, the positively charged Lys were replaced by Asn at 41 and by His at 175 (Fig. S1a). To enhance the binding affinity of PI(4,5)P_2_diC_8_ towards Kir6.2, we made mutations N41K or H175K on Kir6.2. Neomycin is a polyvalent cation that can bind and dissociate PI(4,5)P_2_ from Kir6.2^33^ and high neomycin sensitivity is correlated with low PI(4,5)P_2_ affinity ^33^. Therefore, we exploited the neomycin sensitivity assay to gauge the PI(4,5)P_2_ affinity of Kir6.2 mutants. We found that the H175K mutation significantly reduced the neomycin sensitivity, indicating an enhanced PI(4,5)P_2_ affinity (Fig. 1a). Further analysis showed that H175K mutant can be inhibited by ATP and activated by Mg-ADP and NN414 (Fig. 1b), which is a high-affinity diazoxide-type K_ATP_ opener (Fig. S1b) ^34^.

**Figure 1.**
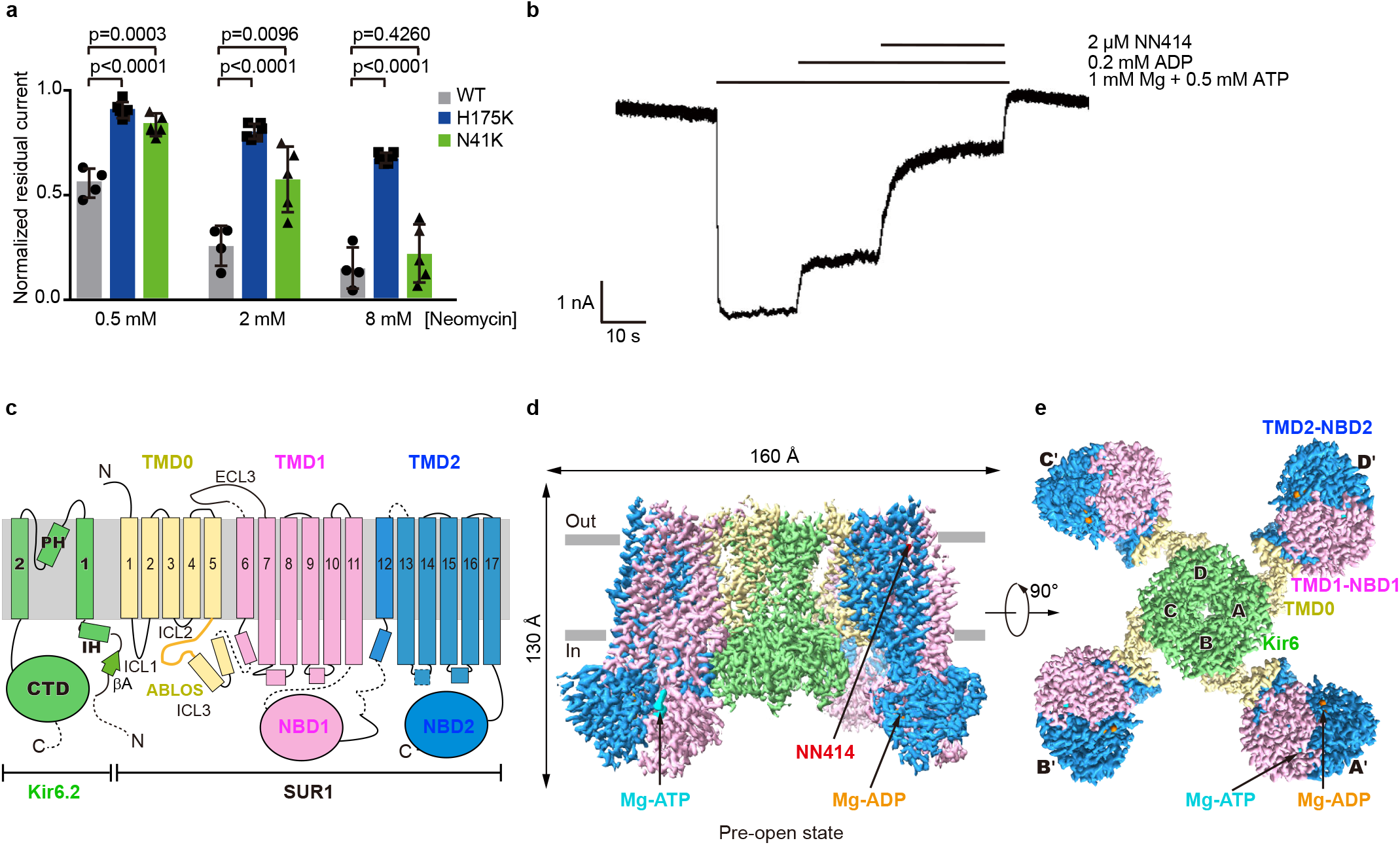
Structure of the pancreatic K_ATP_ channel (H175K_cryoEM_) in the pre-open state. **a,** Neomycin inhibition of the inside-out currents of the K_ATP_ channel. Wild type (WT), H175K, and N41K mutants of Kir6.2 were co-expressed with wild type SUR1 for recordings. Data is shown as mean ± SD, n= 4–6 independent patches. p-value was calculated by unpaired two-tailed t-test. **b,** Representative inside-out recordings of K_ATP_ channel with H175K mutation on Kir6.2. **c,** Topology of Kir6.2 and SUR1 subunits. PH, pore helix; ECL, extracellular loop; ICL, intracellular loop; IH, interfacial helix; CTD, cytoplasmic domain; TMD, transmembrane domain; NBD, nucleotide-binding domain. Transmembrane helices are shown as cylinders. The phospholipid bilayer is shown as thick gray lines. Kir6.2, SUR1 TMD0-ICL3 fragment, TMD1-NBD1, and TMD2-NBD2 are shown in green, yellow, violet, and blue, respectively. **d,** Side view of the K_ATP_ complex in the pre-open state. Mg-ADP, Mg- ATP, and NN414 are shown in orange, cyan, and red, respectively. **e,** Bottom view of the K_ATP_ channel in the pre-open state from the intracellular side.

Based on these observations, we made H175K mutation on the SUR1-Kir6.2 fusion constructs, in which the C terminus of SUR1 is covalently linked to the N terminus of Kir6.2 by a long linker to ensure the correct 4:4 stoichiometry between Kir6.2 and SUR1 ^26^, yielding the H175K_cryo-EM_ construct. The H175K_cryo-EM_ construct can be inhibited by ATP and activated by Mg-ADP and NN414 (Fig. S1c). These results suggest H175K_cryo-EM_ recapitulates the basic electrophysiological properties of wild type K_ATP_ channel and could be used for structural studies. We purified H175K_cryo-EM_ protein in detergent and supplemented Mg-ADP, PI(4,5)P_2_diC_8_, and NN414 into the protein for cryo-EM sample preparation (Fig. S1d,e).

Single particle cryo-EM analysis showed the H175K_cryo-EM_ protein shows the “propeller” shape (Figs. 1c-e, S2 and S3), similar to our previous wild type K_ATP_ protein in a similar condition ^26^. The consensus refinement revealed that the peripheral ABC transporter modules of the SUR1 subunit (TMD1-NBD1-TMD2-NBD2) show motions relative to the central K_ATP_ channel core, consisting of Kir6.2 and SUR1-TMD0 domains. We further exploited symmetry expansion, signal subtraction, and local refinement to improve the resolution of the ABC transporter module to 3.1 Å^26^ (Fig. S2). The focused 3D classification revealed that the core has obvious conformational heterogeneity at the bundle crossings of the Kir6.2 channel, showing a close to open transition at the gate. Subsequent refinement resolved two 3D classes with a closed gate and pre-open gate, the resolution of which reached 3.16 Å and 2.87 Å for Kir6.2 channel respectively (Fig. S2). The maps obtained from local refinement were aligned to consensus maps and summed to yield two composite maps for model building and interpretation (Fig. S2, S3, and Table S1).

### Conformational changes of Kir6.2 TMD during channel opening

The structure of Kir6.2 in the closed state of H175K_cryo-EM_ is highly similar to our previous ATP+RPG state structure (PDB ID: 6JB1)^27^, with root-mean-square deviation of 0.7198 Å (Figs. 2 and S4). Residues on the M2 helix tightly seal the pore at the bundle crossing (Fig. 2c). The side chains of L164 and F168 form the gate, where the radius of the narrowest restriction is below 1 Å (Fig. 2c). In contrast, in the pre-open state structure, the inner part of the M2 helix moves outward (Fig. 2d-e). Particularly, the side chains of L164 and F168 move away from the center, resulting in the dilation of the pore (Fig. 2). The radius of the ion permeation pathway at the bundle crossing of TM now increases to 3 Å (Fig. 2c). The constriction at the cytosolic G-loop gate has a radius of 2.6 Å (Fig. 2c). Although the Kir6.2 channel in the current structure does not allow the passage of fully hydrated potassium ions (with a radius of 3.3 Å), compared to the CNG channel in open state (PDB ID: 6WEK)^35^, the channel begins to open.

**Figure 2.**
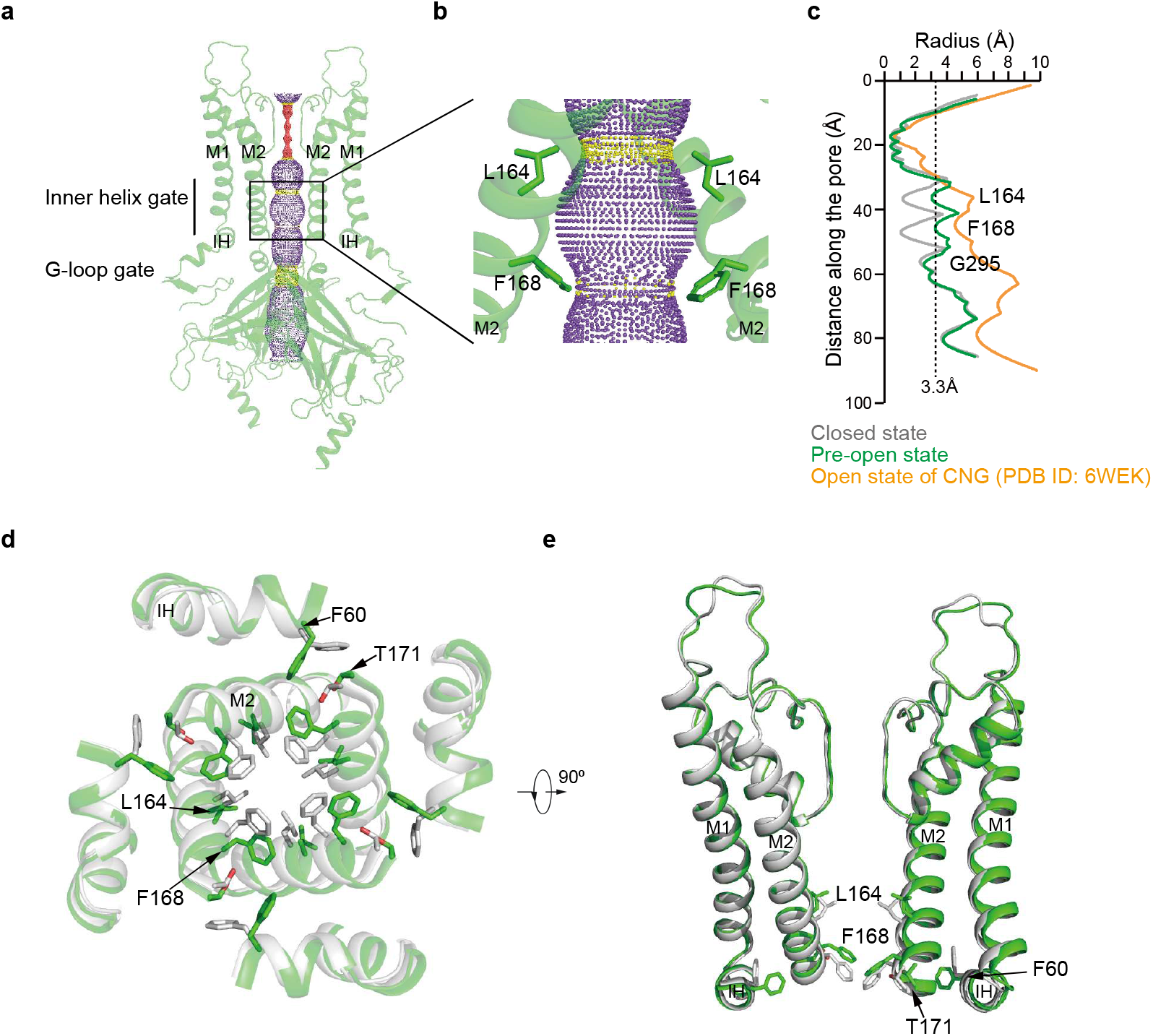
The conformational changes of Kir6.2 TMD during K_ATP_ channel opening. **a,** Side view of Kir6.2 subunits of H175K_cryoEM_ in the pre-open state. The ion conduction pathway along the pore is shown as dots and colored as red, yellow, and purple according to the pore radii of <1.4, 1.4-3.3, and >3.3 Å. M1, M2, and IH are labeled. For clarity, the subunits in front and in the back were omitted. **b,** Close-up view of M2 helices with gate residues shown as sticks (L164 and F168). **c,** Calculated pore profiles of the H175K_cryoEM_ closed state (gray), H175K_cryoEM_ pre-open state (green), and the open state of CNG channel (PDB ID: 6WEK)(orange). The size of a fully hydrated potassium ion (3.3 Å) is shown as dashes. **d,** Structural comparison of the transmembrane domain between the closed state (gray) and the pre-open state (green). **e**, a 90° rotated view of **d**.

Associated with the expansion of the pore at the center, there are concomitant movements of the inner part of the Interfacial Helix (IH) and M1 helices (Fig. 2d). In the closed state, the side chains of F60 on IH pack against T171 on M2 to stabilize the closed pore (Fig. 2d,e). While in the pre-open state, the side chains of F60 swing away, which allows the expansion of M2 (Fig. 2d,e). In contrast to the obvious structural rearrangements of the inner portion of the pore, the structure of outer region of the pore, especially at the selectivity filter, has little change (Fig. 2e).

In the structure of Kir3.2 in complex with PI(4,5)P_2_diC_8_, the PI(4,5)P_2_diC_8_ molecules bind at subunit interface at the inner leaflet of the membrane. We found lipid-like densities in both closed state and pre-open state of H175K_cryo-EM_ at similar positions (Fig. S3), but the lack of head group features hindered confident identification of their identities. Therefore, whether PI(4,5)P_2_diC_8_ molecules were bound in the H175K_cryo-EM_ structures await further investigation.

### Conformational changes of Kir6.2 CTD during channel opening

In the closed state, there are ADP densities inside each nucleotide-binding pocket of Kir6.2 CTD (Fig. S3). The binding mode of the adenosine group of ADP is similar to that observed previously^25–27^. In contrast, there is no ADP density in the pre-open state, and we observed an obvious structural reorganization around the nucleotide-binding pocket of Kir6.2 (Fig. 3). Conformation of R50-R54 on the βA-IH loop has large changes (Fig. 3a,b). The side chains of Q52 flip from solvent-exposed conformation to buried conformation, while the side chains of E51 move in the opposite direction, occupying the nucleotide-binding pocket (Fig. 3b). During channel opening, there is a 6° anti-clockwise rotation of CTD viewing from the intracellular side, resulting in the shrinkage of nucleotide-binding pocket in the pre-open state (Fig. 3c). Therefore, the structure of Kir6.2 in the pre-open state is not compatible with the binding of nucleotides anymore, in agreement with the fact that no ADP was observed in the nucleotide-binding pocket of the pre-open state.

**Figure 3.**
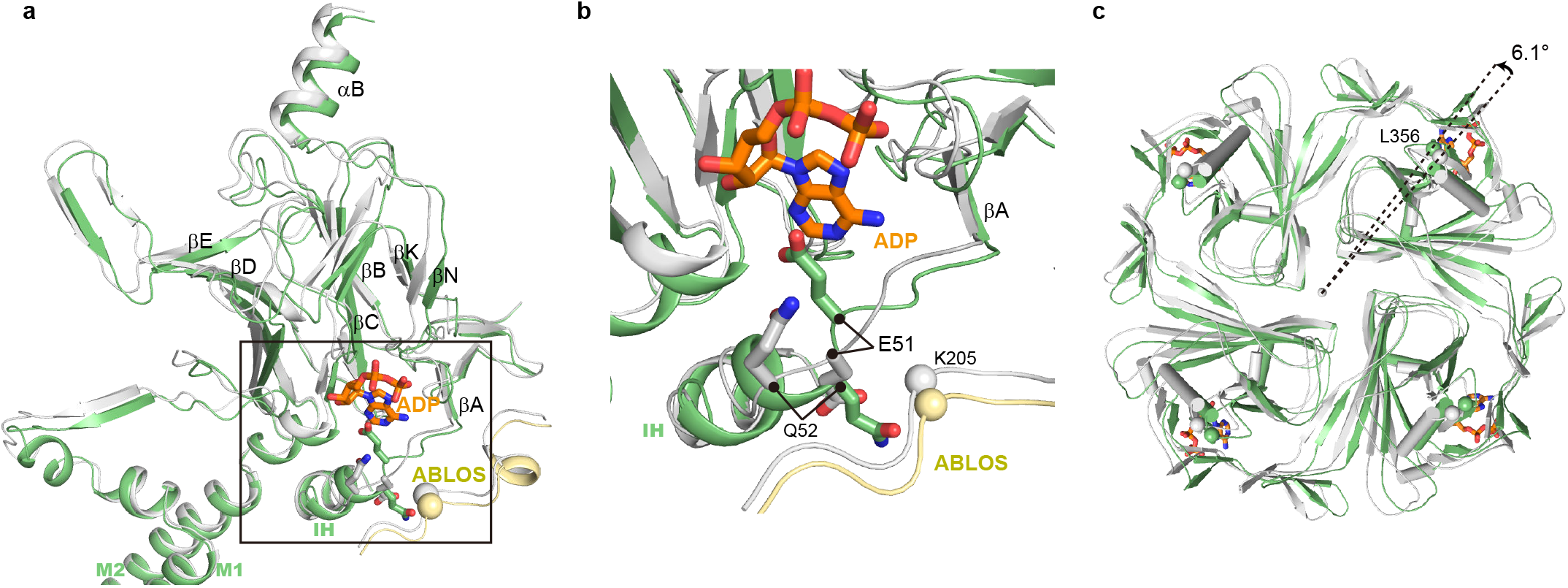
Conformational changes of Kir6.2 CTD during K_ATP_ channel opening. **a,** Conformational changes of the nucleotide-binding site between the closed (gray) and the pre-open (green) states of H175K_cryoEM_. The ADP bound in the closed state is shown as sticks in orange. **b,** Close-up view of the nucleotide-binding site boxed in **a**. **c,** Bottom view of the Kir6.2 CTD. The rotation angle between CTDs was measured using Cα positions of L356 of Kir6.2 as marker atoms (shown as spheres).

### Conformational changes of SUR1 TMD0 domain during channel opening

TMD0 domain of SUR1 has a five-helix-bundle structure ^23, 25^. The N terminal region and M1 of SUR1 TMD0 interact with the M1 helix of Kir6.2. The extracellular side of TMD0 has the docking groove for ECL3 of SUR1 ABC transporter module (335-347) ^27^. We observed the outward movements of the TMD0 domain in the inner leaflet and cytosolic region (ICL1, ICL2, and ICL3) during channel opening, while the structure of TMD0 in the outer leaflet largely stays the same (Fig. 4a). These observations suggest the outer half of TMD0 is a structural scaffold that is responsible for tethering Kir6.2 and the ABC transporter module, while the inner half of TMD0 has conformational plasticity for regulatory function. In detail, part of the ICL1 (51-60) of TMD0 is disordered in the closed state, but it is ordered in the pre-open state and forms main-chain hydrogen bonding with βA of Kir6.2 CTD (Fig. 4a,b). In the RPG+ATP bound state structure (PDB ID: 6JB1), K205 on the ATP-binding loop of SUR1 (ABLOS) motif of ICL3 from TMD0 interacts with β and γ phosphates of ATP . But in both the closed state or pre-open state structures of H175K_cryoEM_, we found K205 is away from ADP and does not form interaction with β phosphate of ADP (Figs. 4a, c, d and S4). Moreover, the side-chain densities of residues on the ABLOS motif are not well resolved, indicating their high mobility. By measuring the distances between marker atoms of adjacent subunits, we found coordinated outward movements of Kir6.2 M1 helices (K67 of Kir6.2), the ABLOS motif (K205 of SUR1) and SUR1 ABC transporter module (K394 of SUR1) (Fig. 4c,d).

**Figure 4.**
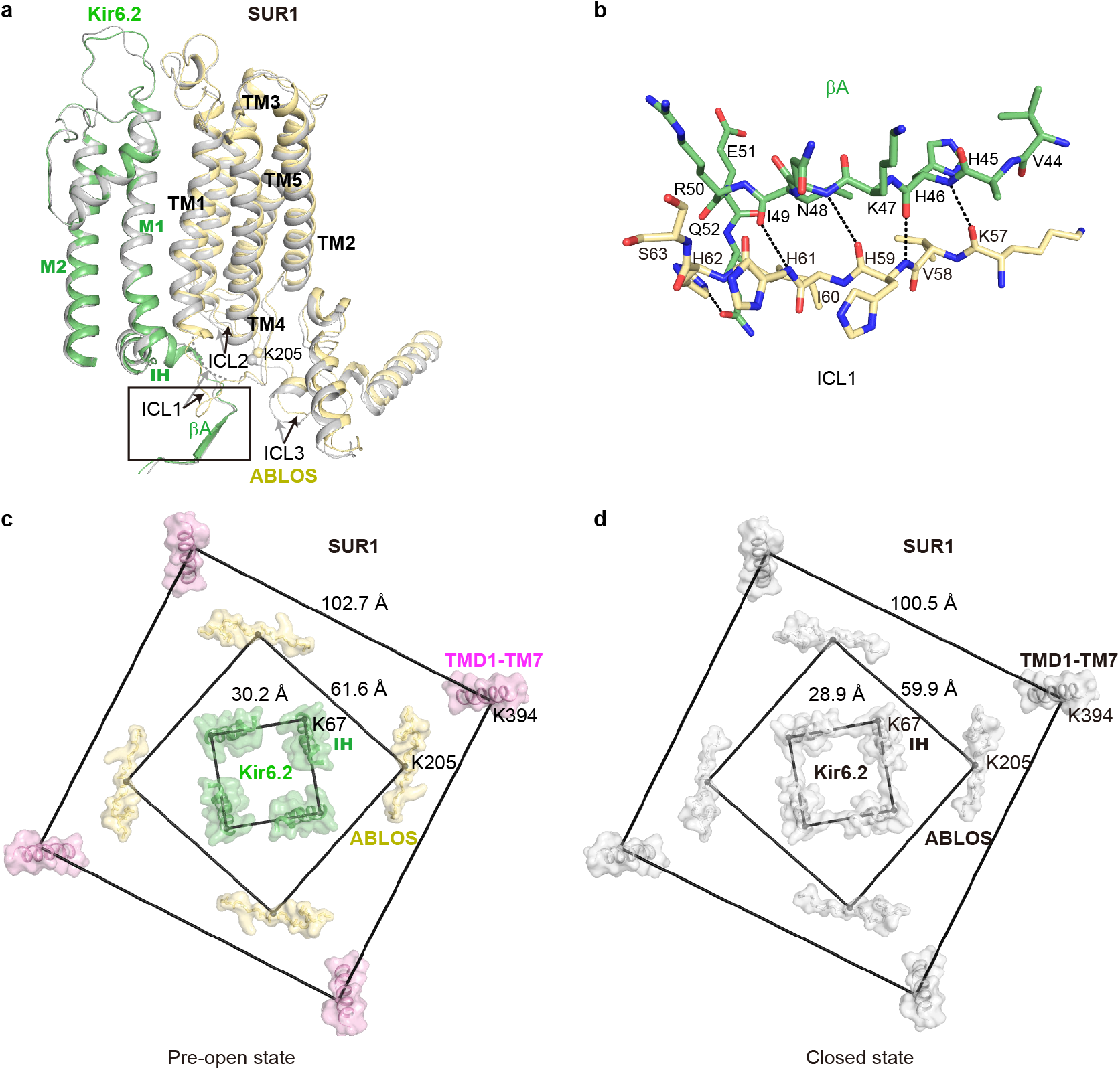
Conformational changes of SUR1 during K_ATP_ channel opening. **a,** Structural comparison of SUR1 TMD0 between the closed state (gray) and the pre-open state (colored) of H175K_cryoEM_. **b,** Close-up view of the interaction between βA of Kir6.2 and SUR1-ICL1 boxed in **a**. Putative hydrogen bondings are shown as dashes. **c-d,** Bottom view of the structural arrangement of the K_ATP_ complex during channel opening. Cα positions of K67 on IH of Kir6.2, K205 on ABLOS of SUR1, and K397 on M7 of SUR1 are shown as spheres. Distances of marker atoms in the pre-open state (colored) and the closed state (gray) are shown in **c** and **d**, respectively.

### KCO binds inside the SUR1 subunit in the NBD-dimerized conformation

The ABC transporter module of SUR1 shows an asymmetric NBD-dimerized structure as observed previously^22, 26^ (Fig. 5a). We observed that Mg-ADP is bound in the partially closed consensus site, while Mg-ATP is bound in the fully closed degenerate site (Fig. 5b,c). Since we did not supplement ATP into the cryo-EM sample, the ATP molecules might be carried through purification. The excellent local map quality allowed us to unambiguously identify the NN414 molecule and determine its binding pose (Fig. 5d), which was further confirmed with the aid of computational methods (see Methods section).

**Figure 5.**
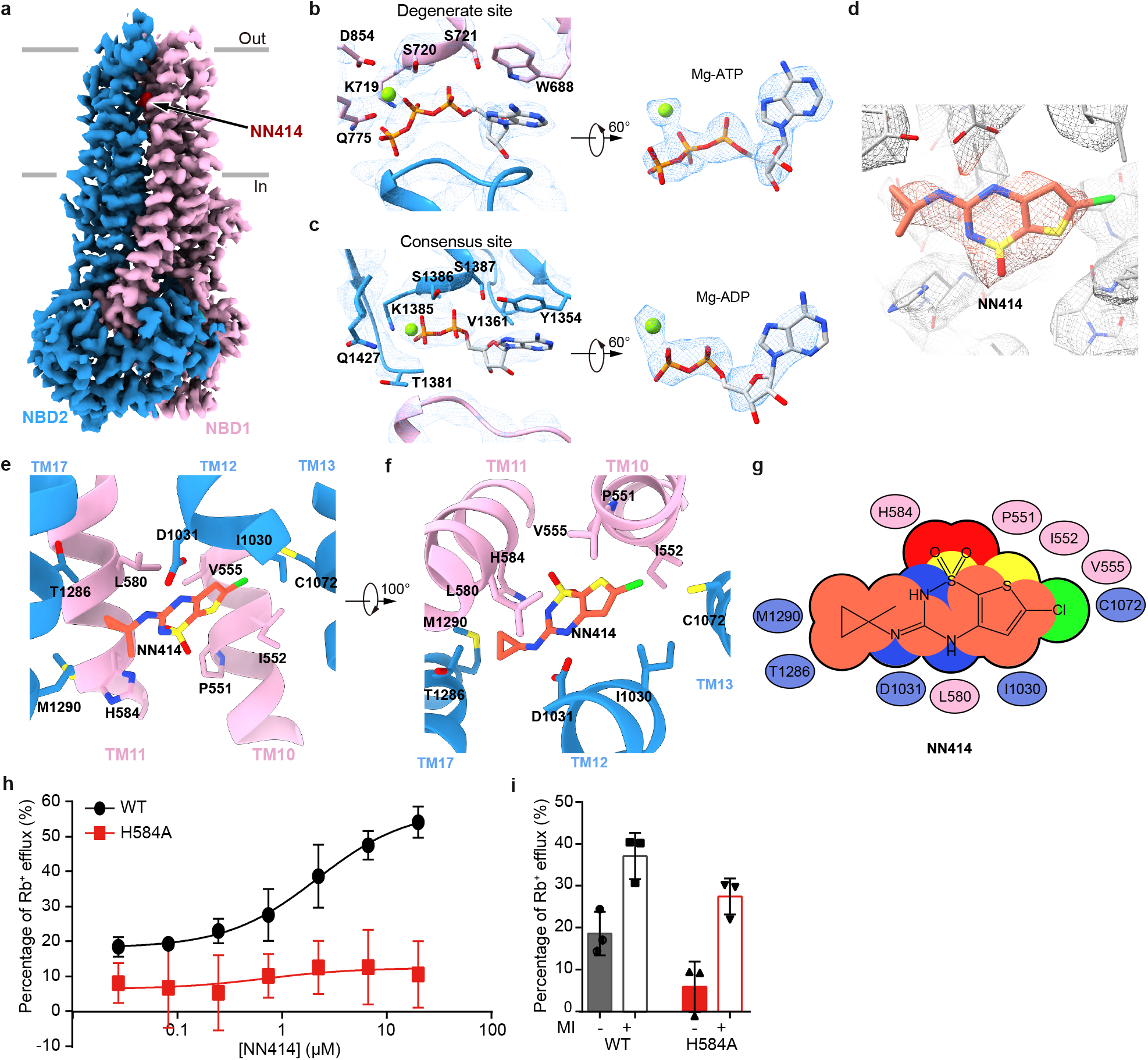
Structure of the SUR1 subunit in complex with NN414 and Mg-nucleotides. **a,** Cryo-EM density map of SUR1 in complex with Mg-nucleotides and NN414, viewed from the side. The approximate position of the lipid bilayer is indicated by gray bars. TMD1- NBD1, TMD2-NBD2, and NN414 are colored in pink, blue and red, respectively. For better visualization of the position of NN414, a fragment of TMD2 in front of NN414 was omitted. **b,** Close-up views of electron densities at the degenerate site. NBD1, NBD2, nucleotides, and Mg^2+^ are colored in pink, blue, gray, and green, respectively. **c**, Electron densities at the consensus sites. **d,** NN414 density (orange) in the SUR1 subunit (gray). The map is shown as mesh and the protein is shown as sticks. **e–f,** Close-up views of the NN414-binding site. TMD1 and TMD2 are colored in pink and blue, respectively. NN414 (orange) and residues that interact with NN414 are shown as sticks. **g,** Cartoon representation of the interaction between NN414 and SUR1. The key residues on TMD1 and TMD2 are shown as pink and blue ovals, respectively. **h**, The dose-response activation curves of SUR1- Kir6.2 K_ATP_ channel by NN414 measured by Rb^+^ efflux assay. Curves were fitted to the Hill equation. Error bars represent the standard error and n≥3. **i**, Effects of metabolism inhibitors on K_ATP_ channel containing various SUR1 mutants. Error bars represent the standard error and n≥3.

NN414 binds in the KCOS of the SUR1 subunit. The dioxide group and its adjacent nitrogen of NN414 form polar interactions with H584 on TM11 (Fig. 5e-g). One NH group on the benzothiadiazine ring and the other NH group between benzothiadiazine ring and methylcyclopropyl group of NN414 form hydrogen bonding with D1031 on M12 (Fig. 5e-g). Several hydrophobic interactions further stabilize NN414 binding. The methylcyclopropyl group of NN414 forms hydrophobic interactions with M1290, Y1287, and T1286 on M17 (Fig. 5e-g). The central benzothiadiazine ring of NN414 is sandwiched by C1072 of M13, L1030 of M12, and I552 of M10 on one side and V555 of M10 and L580 of M11 on the other side (Fig. 5e-g). Rb^+^ efflux assay showed that H584A mutation does not affect K_ATP_ channel activation induced by metabolic inhibitors but abolished the activation by NN414, suggesting its essential role in the binding of NN414 (Fig. 5h-i).

## Discussions

### Relationship between Kir6.2 CTD rotation and channel opening

In our previous structures of pancreatic K_ATP_ in the presence of ATP+RPG^27^, ATP+GBM, or Mg-ADP+NN414^26^, we all observed the “propeller” structures but their CTDs show both “R” state and “T” state, with 12-13° anticlockwise rotation from “R” state to “T” state, viewed from the cytosolic side. These structures are similar to the Kir3.2 structure which shows “undocked state” (similar to “R” state) in the absence of PI(4,5)P_2_ or docked state (similar to “T” state) in the presence of PI(4,5)P_2_ ^32^. In the current work, we present the structures of H175K_cryoEM_ at both closed state and pre-open state. The CTD of H175K_cryoEM_ in closed state is in a similar position to the “T” state of ATP+RPG structure (PDB ID: 6JB1) (Fig. S4c). While the CTD of the pre-open structure has additional 6° anti-clockwise rotation compared to the “T” state (Fig. 3c). These structural observations support the idea that K_ATP_ opening is associated with the rotation of Kir6.2 CTD, similar to that proposed for Kir2.2 ^31^ or Kir3.2 channel^32^, indicating a conserved “rotate to open” mechanism for Kir channel gating.

### Mechanism of nucleotide inhibition of K_ATP_ channel

The cryo-EM density maps showed that in the same sample, inhibitory ADP molecules exclusively bind Kir6.2 in the closed state but not in the open state. Further structural analysis revealed that the conformational changes of Kir6.2 during channel opening, including rotation of CTD and structural rearrangement of the βA-IH loop, disrupt the inhibitory nucleotide-binding pocket of Kir6.2 (Fig. 3a,b). Conversely, the wedging of nucleotide inside the nucleotide-binding pocket of the Kir6.2 channel would block conformational changes that are required for channel opening, providing a plausible mechanism for Kir6.2 channel inhibition by nucleotides. The signals of nucleotide binding in Kir6.2 CTD are further transmitted to the central ion channel pore via several structural elements, including CTD, βA-IH loop, IH, and M2 gating helix (Fig. 3a). Corroborating with our observations, these structural elements are hotspots for genetic mutation of neonatal diabetes outside the ATP binding pocket, such as E51, Q52, and G53 on βA-IH linker; V59, F60, and V64 on IH helix; A161, L164, C166, I167 and K170 on M2(Fig. S5) ^5^, suggesting that mutations in these structural elements might allosterically affect ATP inhibition and channel gating.

### Mechanism of Mg-ADP and KCO activation of K_ATP_ channel

The binding of Mg-ADP to SUR1 induces the asymmetric dimerization of NBD1 and NBD2, which further drives the closure between TMD1 and TMD2 of the SUR1 ABC transporter module. Our structure shows that diazoxide-type K_ATP_ openers, exemplified by NN414, interact with both TMD1 and TMD2 (Fig. 5), to promote the closure of TMD (Fig. 6). The converged structural changes induced by Mg-ADP and K_ATP_ openers suggest their synergistic effect on K_ATP_ activation.

**Figure 6.**
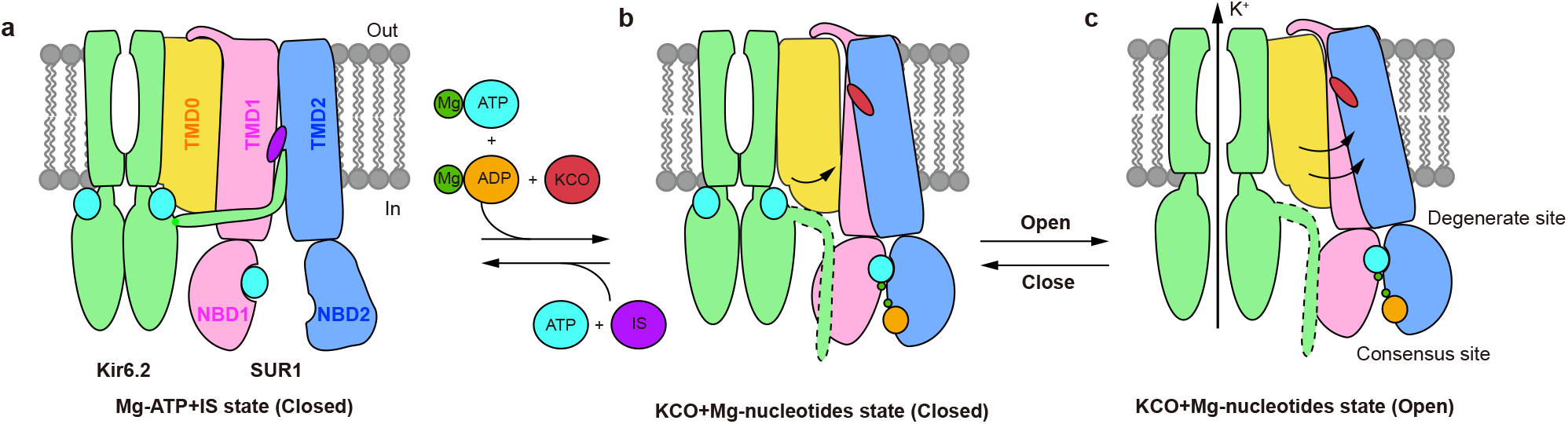
Model for the K_ATP_ channel activation by Mg-nucleotides and K_ATP_ opener. **a– c,** Side view of the cartoon model of the K_ATP_ channel. For simplicity, a pair of Kir6 subunits and one SUR2 subunit are shown. Kir6, TMD0, TMD1-NBD1, TMD2-NBD2, Mg^2+^, ATP, ADP, insulin secretagogue (IS), and K_ATP_ opener (KCO) are colored in green, yellow, pink, blue, dark green, cyan, orange, purple, and red, respectively. The flexible KNtp in the Mg- nucleotide and K_ATP_ opener-bound state is outlined by dashed lines.

Comparing the structure of K_ATP_ in the presence of ATP+RPG (PDB ID: 6JB1) ^27^ with the structure of H175K_cryoEM_ in the closed state (Fig. S4a,b), we found that their Kir6.2 channels are all in the same nucleotide-bound inhibited conformation (Fig. S4c). In contrast, there is a large structural rearrangement of SUR1 due to Mg-ADP and NN414 binding (Fig. S4d-f). The conformational changes of the ABC transporter module are transmitted to the lasso motif and finally arrive at TMD0, resulting in outward tilting of the inner half of TMD0 (Fig. S4d-f). Notably, the important ATP-coordinating residue K205 on the ABLOS motif of SUR1 ^27^ moves outward and is away from the inhibitory nucleotide bound on the Kir6.2 (Fig. S4d-f). This would certainly weaken the binding of inhibitory nucleotides and in turn, promote channel activation (Fig. 6a,b). During the opening of Kir6.2, there is a further outward tilting of TMD0 and its associated intracellular loops, leaving more space for the expansion of Kir6.2 TMD (Figs. 4 and 6b,c). Together with outward movement of ABLOS motif on SUR1 ICL3 where E203 locates (Fig. 4), there is a dramatic flipping movement of Q52 on βA-IH of Kir6.2 (Fig. 3). It is reported that Q52E mutation of Kir6.2 paired with E203K of SUR1 mutation greatly enhances the ATP sensitivity of K_ATP_, and oxidative crosslinking of Q52C (Kir6.2) and E203C (SUR1) mutant could lock the channel in a closed state ^36^. Our structural observation suggests that these two mutations, on one hand, block the opening conformational change of Kir6.2 directly and, on the other hand, fix the relative distance between the SUR1 ABLOS motif and Kir6.2 to inhibit channel opening.

Kir6 N-terminal peptide (KNtp) plays important role in regulating K_ATP_ function^29^. KNtp not only enhances the ATP sensitivity of Kir6.2 but also mediates the inhibition of insulin secretagogue in the absence of nucleotides, possibly by binding to the central vestibule of SUR and restraining the mobility of Kir6 CTD (Fig. 6a) ^29, 30^. In the H175K_cryo-EM_ construct, the KNtp is covalently fused to the C-terminus of SUR1 and therefore could not bind inside the SUR1 central vestibule anymore ^26, 27^. However, the H175K_cryo-EM_ construct could be inhibited by ATP and activated by Mg-ADP, as wild type K_ATP_ channel (Fig. S1c). Therefore, our work current revealed the KNtp-independent nucleotide regulation mechanism of K_ATP_ channels (Fig. 6). The role of KNtp during K_ATP_ channel opening awaits further studies.

## Materials and Methods

### Cell lines

FreeStyle 293-F (Thermo Fisher Scientific) suspension cells were cultured in SMM 293-TI (Sino Biological Inc.) supplemented with 1% FBS at 37 ℃, with 6% CO2 and 70% humidity. Sf9 insect (Thermo Fisher Scientific) cells were cultured in Sf-900 III SFM medium (Thermo Fisher Scientific) at 27℃. AD293 cells (Agilent) were cultured in DMEM basic (Thermo Fisher Scientific) supplemented with 10% fetal bovine serum (FBS) at 37 ℃, with 6% CO_2_ and 70% humidity.

### Method Details

#### Construct of H175K_cyro-EM_

We used cDNA of SUR1 from *Mesocricetus auratus* (maSUR1) *and cDNA of Kir6.2 from Mus musculus* (mmKir6.2) for our studies. We made a maSUR1-mmKir6.2 H175K fusion construct (H175K_cryoEM_) which is similar to our previous K_ATP_ fusion construct^26^. There were a 41-residue linker (VDGSGSGSGSAAGSGSGSGSGSGAAGSGSGSGSGSGAAALE) and an 8-residue Prescission Protease cleavage site (LEVLFQGP) between SUR1 and Kir6.2 H175K mutant. The first Met of Kir6.2 was removed to minimize internal translation initiation. This construct contains C-terminal GFP tag and strep tag which were used for protein purification. For electrophysiological experiments, Kir6.2 was cloned into modified C-terminal GFP-tagged BacMam expression vector and SUR1 into non-tag BacMam expression vector as described previously{Wu, 2018 #646} .

### Electrophysiology

K_ATP_ constructs were transfected into FreeStyle 293-F cells using polyethylenimine at the cell density of 0.8∼1.1×10^6^ cells/ml. Cells were cultured in 293TI medium supplemented with FBS for 24∼36 hr before recording. Macroscopic currents were recorded in inside-out mode at +60 mV via Axon-patch 200B amplifier (Axon Instruments, USA). Patch electrodes were pulled by a horizontal microelectrode puller (P-1000, Sutter Instrument Co, USA) to 3.0∼5.0 MΩ resistance. Both pipette and bath solution were KINT buffer, containing (mM): 140 KCl, 1 EGTA and 10 HEPES (pH 7.4, KOH). For neomycin inhibition, the 50 mM stock neomycin (Sigma) was made in DMSO, stored at -20℃and diluted into KINT buffer to a desired concentration. For NN414 activation, the 50 mM NN414 stock (Sigma) was dissolved in DMSO, stored at -20℃ and diluted into KINT buffer to final working concentration. ATP and ADP stocks were prepared on ice, aliquoted and stored at -20℃. ATP and ADP were dissolved in water and adjusted to pH = 7 by KOH. The nucleotide concentration was determined by its extinction coefficient and the UV-absorption at 259 nm. Recordings were acquired at 5 kHz and low-pass filtered at 1 kHz. Data were further analyzed by pClampfit 10.0 software.

### Rb^+^ efflux assay

AD293 cells were cultured in 6-well plate till 90-95% confluence. Wild type mmKir6.2 with C terminal GFP tag was co-transfected with wild type maSUR1 or maSUR1 with H584A mutation into AD293 cells. Cells were continually cultured for 24 hr for protein expression and assembly, and GFP signal was detected by microscope in *vivo*. Then cells were digested by trypsin supplemented with 0.25 mM EDTA and were equably separated into 96-well plate, finally incubated with 100 ul medium per well. For Rb^+^ efflux determination, 6 mM Rb^+^ was supplemented into the medium for Rb^+^ pre-incubation into the cells. The 96-well plate was pre-treated by polylysine for about 24 hr in 37℃, and washed out by DMEM medium without FBS before incubation cells. After incubation in 96-well plate for 12∼14 hr, cells were washed by Ringer’s solution (mM) : 118 NaCl, 10 HEPES (pH 7.4), 25 NaHCO_3_, 4.7 KCl, 1.2 KH_2_PO_4_, 2.5 CaCl_2_, and 1.2 MgSO_4_ and incubated with Ringer’s solution supplemented with NN414 in different concentration for 10 min. As a baseline control, plasmids expressing GFP were transfected into cells at the same time. To detect whether K_ATP_ channels were functionally expressed, metabolic inhibitors including 3mM de-O-glucose and 1 μM oligomycin were used to activate K_ATP_ channels. After drug treatment, the supernatants were transferred for Rb^+^ efflux determination. The cells plated in the well were dissolved by 1% triton X-100 for 30 min and also transferred for total Rb^+^ quantification. The quantification of Rb^+^ was carried out on Ion Channel Reader 8000 (Aurora Group Company).

### Protein expression and purification

K_ATP_ channels were expressed as described previously and the purification process was carried out with minor modification{Wu, 2018 #646}. For protein purification, membrane pellets were homogenized in TBS (20 mM Tris-HCl pH 7.5, 150 mM NaCl) and solubilized in TBS with 1% digitonin (biosyth), supplemented with protease inhibitors (1 mg/ml Leupepetin, 1 mg/ml Pepstatin, 1 mg/ml Aprotonin, and 1 mM PMSF), 1mM EDTA and1mM EGTA for 30 min at 4℃. Unsolubilized materials were removed after centrifugation at 100,000 g for 30 min and supernatant was loaded onto two 5 mL column packed with Streptactin 4FF resin (Smart Lifesciences). Strep column was first washed by TBS buffer supplemented with 0.1% digitonin, protease inhibitors (1 mg/ml leupepetin, 1 mg/ml pepstatin, 1 mg/ml aprotonin), 1mM EDTA and 1mM EGTA. Then the columns were washed with TBS supplemented with 0.1% digitonin, 3mM MgCl_2_ and 1mM ATP. The last washing step buffer was TBS supplemented with 0.1% digitonin. The K_ATP_ octamers were eluted by TBS (50 mM Tris-HCl pH 7.5, 150 mM NaCl) supplemented with 0.1% digitonin and 8 mM desthiobiotin. The eluate was concentrated and loaded onto Superose 6 increase column (GE Healthcare) running with TBS supplemented with 0.1% digitonin. Peak fractions were collected and concentrated to A_280_=15 (estimated as 15 µM K_ATP_ octamers). The protein purification was completed within 15 h and the purified protein was instantly used for cryo-EM sample preparation.

### Cryo-EM sample preparation

K_ATP_ octamers were supplemented with 5 mM MgCl_2_, 0.5 mM ADP, 0.5 mM NN414 and 10 μM PI(4,5)P_2_diC_8_. Acidic PI(4,5)P_2_diC_8_ (Echelon Biosciences) was dissolved in water and titrated by Tris-HCl (pH 7.5) to adjust the pH before usage. Cryo-EM grids were prepared with Vitrobot Mark IV (FEI) and GIG R1/1 holey carbon grids, which were glow-discharged for 120s using air before making Cryo-EM sample grids. 2.5 ul K_ATP_ octamers sample was applied to glow-discharged grid and then the grid was blotted at blotting force in level 2 for 2 s at 100% humidity and 20℃, before plunge-frozen into the liquid ethane.

### Cryo-EM data acquisition

Cryo-grids were screened on Talos Arctica microscope (Thermo Fisher Scientific) operated at 200 kV for small scale data collection. For grids in high quality, a large data set for K_ATP_ channel structure determination was collected in Titan Krios microscope (Thermo Fisher Scientific) operated at 300 kV.

Images were collected using K2 camera (Gatan) which was mounted post a Quantum energy filter with 20 eV slit, operated under super resolution mode with pixel size of 1.324 Å at the object plane, and controlled by Serial EM. Defocus values were set to range from -1.3 μm to - 1.8 μm for data collection. Dose rate on detector was 8 e^-^ s^-1^ A^-2^. And the total exposure was 50 e^-^/A^2^. Each 12 s movie was dose-fractioned into 50 frames.

### Image processing

Movies collected were exposure-filtered, gain-corrected, motion-corrected, mag-distortion-corrected and binned with MotionCor2 ^37^, producing dose-weighted and summed micrographs with pixel size 1.324 Å. CTF models of dose-weighted micrographs were determined using Gctf ^38^. Gautomatch (developed by Kai Zhang, MRC-LMB) was used for particles auto-picking and Gautomatch templates were produced by projecting K_ATP_ density map generated from small scale data collected from 200kV microscope. Data processing was initially executed in Relion_3.1^39^. Particles were extracted from dose-weighted micrographs. After 2 rounds of 2D classification and 3D classification with C4 symmetry, particles of good quality were re-extracted and re-centered. Remaining particles were used for 3D refinement and CTF refinement. Upon convergence, the particles were expanded using C4 symmetry and signals for SUR1 ABC transporter module were subtracted. The subtracted particles were refined using local search within 5° range. The refined SUR1 ABC transporter particles were subjected to no alignment 3D classification, with K=4 and T=20. The 3D classes with good map quality were selected and refined in cryoSPARC2 by non-uniform refinement^40^, CTF refinement and local non-uniform refinement to reach a resolution of 2.7 Å (map-A). Focused 3D classification on Kir6.2 CTD was carried out with K=4 and T=20 without alignment. Two 3D class with good features but with relative rotations were selected and refined using non-uniform refinement, CTF refinement and local non-uniform refinement in cryoSPARC2 to generate consensus maps. Examination of their pore domain revealed they represent the pre-open state and the closed state respectively. With the mask of Kir6.2 for local refinement, the resolution of pre-open state reached 2.87 Å (map-B) and the closed state reached 3.16 Å (map-C). With the mask for Kir6.2 TMD and SUR1 TMD0, the resolution of pre-open state reached 2.94 Å (map-D) and resolution of the closed state reached 3.19 Å (map-E). The sharpened local refined maps were aligned to the consensus map and summed by vop maximum command in chimera to generate composite maps. Specifically, we summed map- A, map-B and map-D to generate pre-open state map and map-A, map-C and map-E to generate closed state map. The composite cryo-EM maps were reboxed to 180×180×180 and used for interpretation, model building, refinement and illustration.

### Model building

The structure of K_ATP_ in complex with ATP and RPG (PDB ID: 6JB1)^27^ or Mg-ADP and NN414 (PDB ID: 5YWC) ^26^ was divided into individual domains and fitted into the cryo-EM maps using UCSF chimera^41^. The model was further manually rebuilt and refined against the maps using Phenix^42^. Figures were prepared with Pymol, UCSF chimera or UCSF Chimera X^43^. Permeation pathways were calculated using HOLE2^44^.

### Computational methods to optimize ligand binding pose

We used LeDock software ^45^ to predict the binding modes of NN414. We completed the protein model using the LePro module (with the default values) and the NN414 model with Rdkit (from open-source cheminformatics; https://www.rdkit.org). The docking was run with the default parameters provided by the LeDock. We chose two top binding poses for future evaluation and analysis. To find which pose of NN414 has the minimum energy, we ran MD (Molecular Dynamics) simulations to evaluate the stability of the docking poses. The MD simulations were performed on graphics processor units (GPUs) with AMBER20’s GPU-accelerated PMEMD simulation code ^46^. All MD simulations used the AMBER ff14SB Force field ^47^ for the protein and gaff2 for ligand. The complex systems were placed in a box of TIP3P water model ^48^ and simulated at a constant temperature of 298.15K. All bonds containing hydrogens were constrained with the SHAKE algorithm. We also calculated the MM/GBSA (Molecular Mechanics/Generalized Born Surface Area) to estimate the binding energy. We simulated 10 ns for MM/GBSA. Further, we performed a long MD simulation to evaluate system stability.

Relative binding free energy (RBFE) calculations are the most common and widespread applications of FEP for drug discovery projects. We used the XFEP platform which have been developed by XtalPi for the FEP calculations ^49^ The XFEP platform constitutes the backbone of an integrated workflow coupling FEP calculations with AI models and wet-lab experiments. XFEP utilizes the AMBER software package ^46^ for free energy calculations. All simulations were performed using the TIP3P water model, as well as AMBER ff14SB for the proteins and XForce Field (XFF) ^49^, developed by XtalPi, with a system-specific force field refinement protocol for the ligands. All simulations used a Langevin integrator with a 2 fs timestep for heating and equilibration, 4 fs for production, and a friction coefficient of 2 ps^−1^. In this work, the simulation time for each λ window was 2 ns. Simulations were repeated three times.

### Quantification and statistical analysis

Global resolution estimations of cryo-EM density maps are based on the 0.143 Fourier Shell Correlation criterion ^50^. The local resolution map was calculated using cryoSPARC ^51^. Rb^+^ efflux assay curves were fitted to the Hill equation using GraphPad Prism 6. Electrophysiological data reported were analyzed with pclampfit 10.0 software. The number of biological replicates (N) and the relevant statistical parameters for each experiment (such as mean or standard error) are described in figure legends. No statistical methods were used to pre-determine sample sizes.

### Materials Availability

Reagents generated in this study will be made available on request, but we may require a payment and/or a completed Materials Transfer Agreement if there is potential for commercial application.

### Data and Code Availability

Atomic coordinates and cryo-EM maps are deposited in EMDB and PDB as follows: pre-open state: EMD-32310 and PDB 7W4O; closed state: EMD-32311 and PDB 7W4P.

## Acknowledgments

We thank Yaxiong Yang, and Tianyi Hou for their kindly help with illustration. We thank Joseph Bryan for sharing maSUR1 cDNAs and J. Marc Simard for sharing mmKir6.2 cDNA. We thank Guifang Duan and Prof. Zhuo Huang in the State Key Laboratory of Natural and Biomimetic Drugs, School of Pharmaceutical Sciences, Peking University for assistance of Rb^+^ efflux assay. Cryo-EM data collection was supported by Electron microscopy laboratory and Cryo-EM platform of Peking University with assistance of Xuemei Li, Zhenxi Guo, Bo Shao, Xia Pei and Guopeng Wang. Part of the structural computation was also performed on the Computing Platform of the Center for Life Science and High-performance Computing Platform of Peking University. We thank the National Center for Protein Sciences at Peking University in Beijing, China for assistance with negative stain EM. The work is supported by grants from National Natural Science Foundation of China (91957201, 31870833 and 31821091 L.C.) This work is also partially supported by Li Ge-Zhao Ning Life Science Young Scholar grant of Peking University.

## Author contributions

Lei Chen initiated the project. Mengmeng Wang purified protein, prepared the cryo-EM sample, and performed electrophysiology experiments. Dian Ding made mutants for SUR1. Mengmeng Wang and Jing-Xiang Wu collected the data. Mengmeng Wang processed the data, built and refined the model with the help of Lei Chen. Xinli Duan, Songling Ma, and Lipeng Lai performed the computational analysis of drug binding pose. All authors contributed to the manuscript preparation.

---

**Figure S1.**
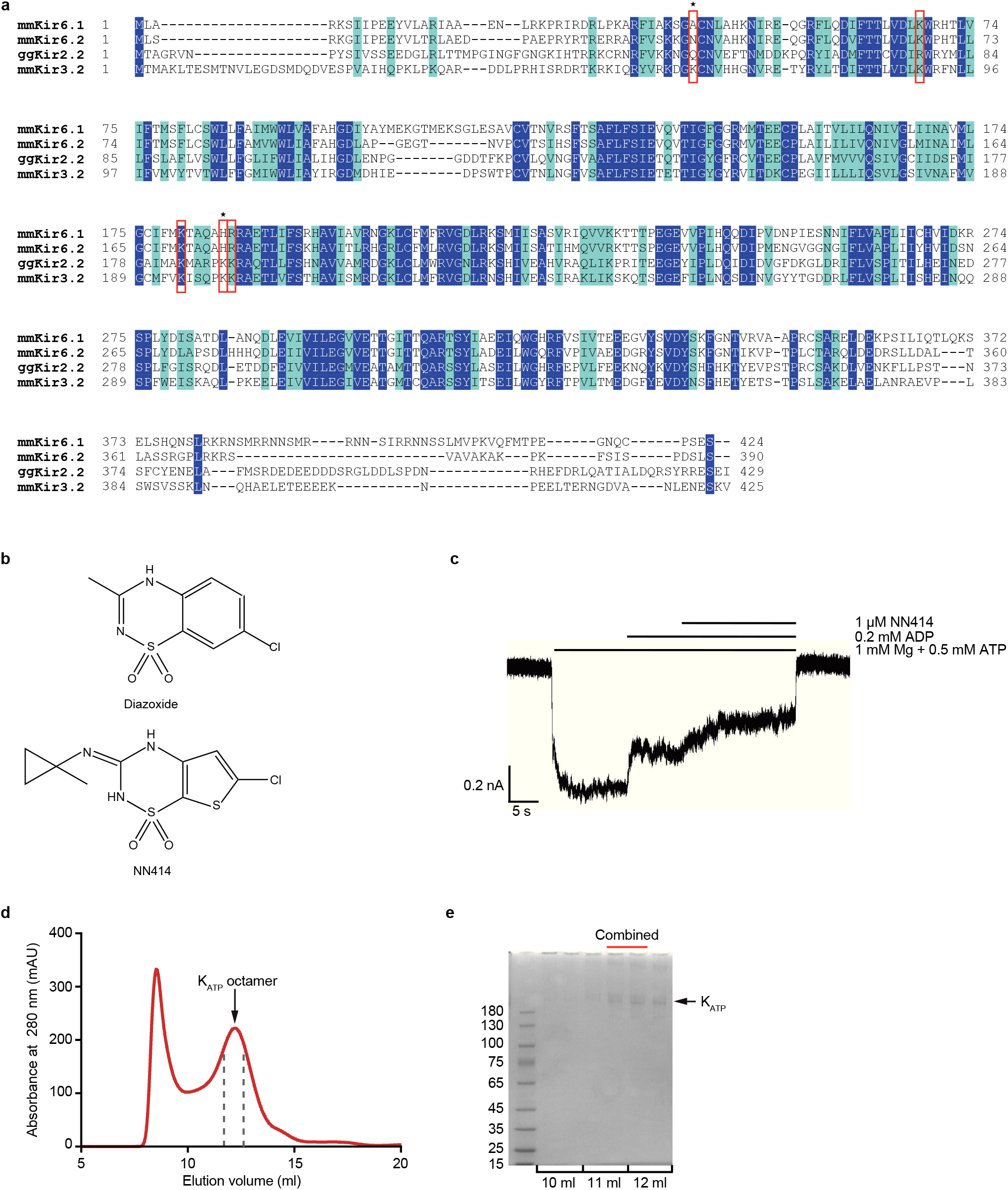
Characterization of K_ATP_ H175K_cryoEM_. **a,** Sequence alignment of Kir6.1 from *Mus musculus* (mmKir6.1), mmKir6.2, Kir2.2 from *Gallus gallus* (ggKir2.2), and mmKir3.2. Identical residues are in the blue background while residues with similarity more than 50% are in cyan background. Positively charged residues that might participate in PI(4,5)P_2_ binding were boxed in red. N41 and H175 of mmKir6.2 were labeled by asterisks above. **b**, Chemical structures of NN414 and diazoxide show their similarities. **c**, Representative inside-out current of H175K_cryo-EM_. **d**, Size-exclusion chromatography (SEC) elution profile of H175K_cryo-EM._ Fractions between the dashed lines were K_ATP_ octamer used for cryoEM sample preparation. **e**, SDS-PAGE of H175K_cryoEM._ Fractions labeled by red bars were collected for cryoEM sample preparation.

**Figure S2.**
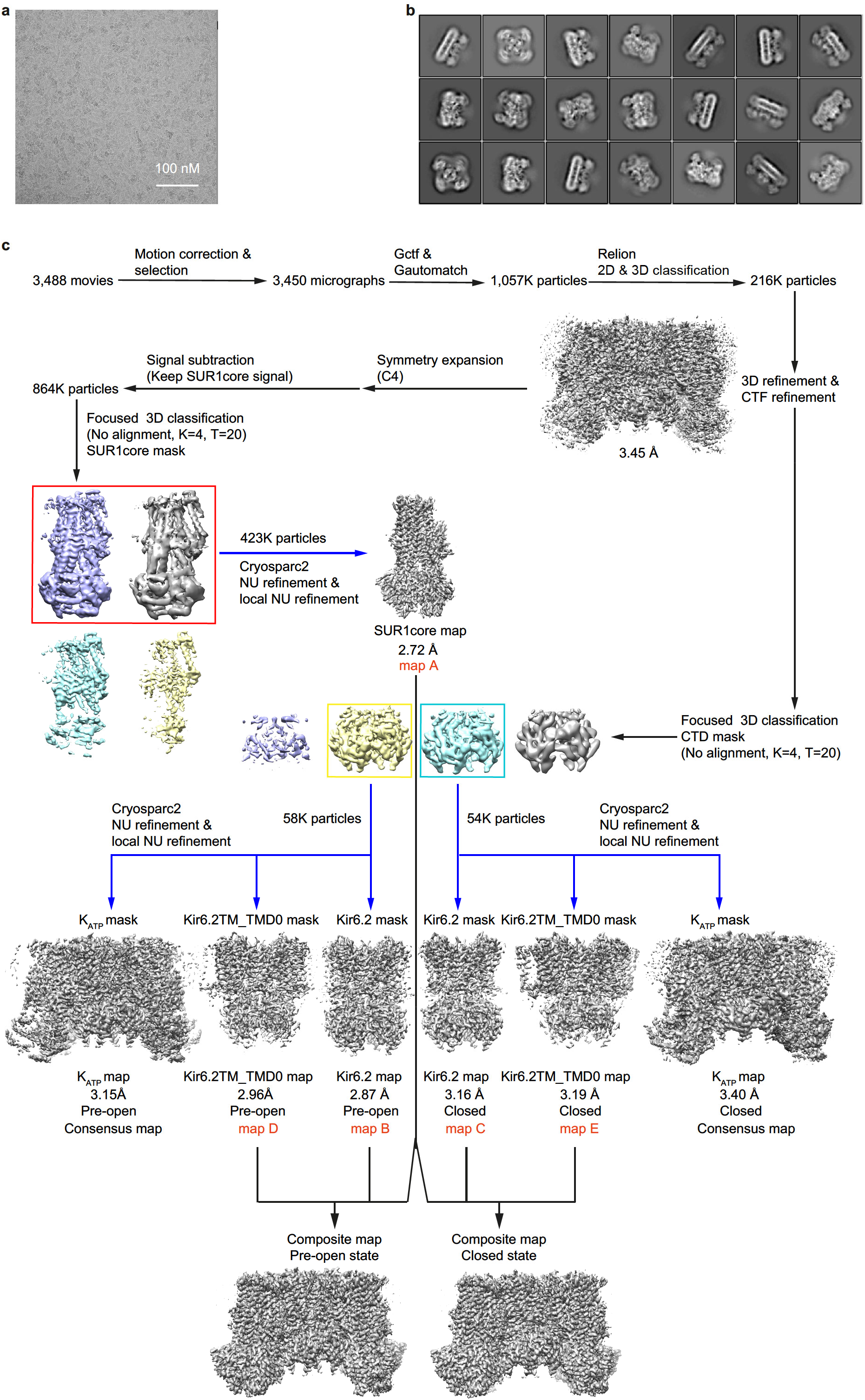
Cryo-EM image processing flow-chart. **a,** Representative raw micrograph. **b,** Representative 2D class averages of H175K_cryo-EM_. **c,** Flow-chart of cryo-EM image processing for H175K_cryo-EM_ dataset.

**Figure S3.**
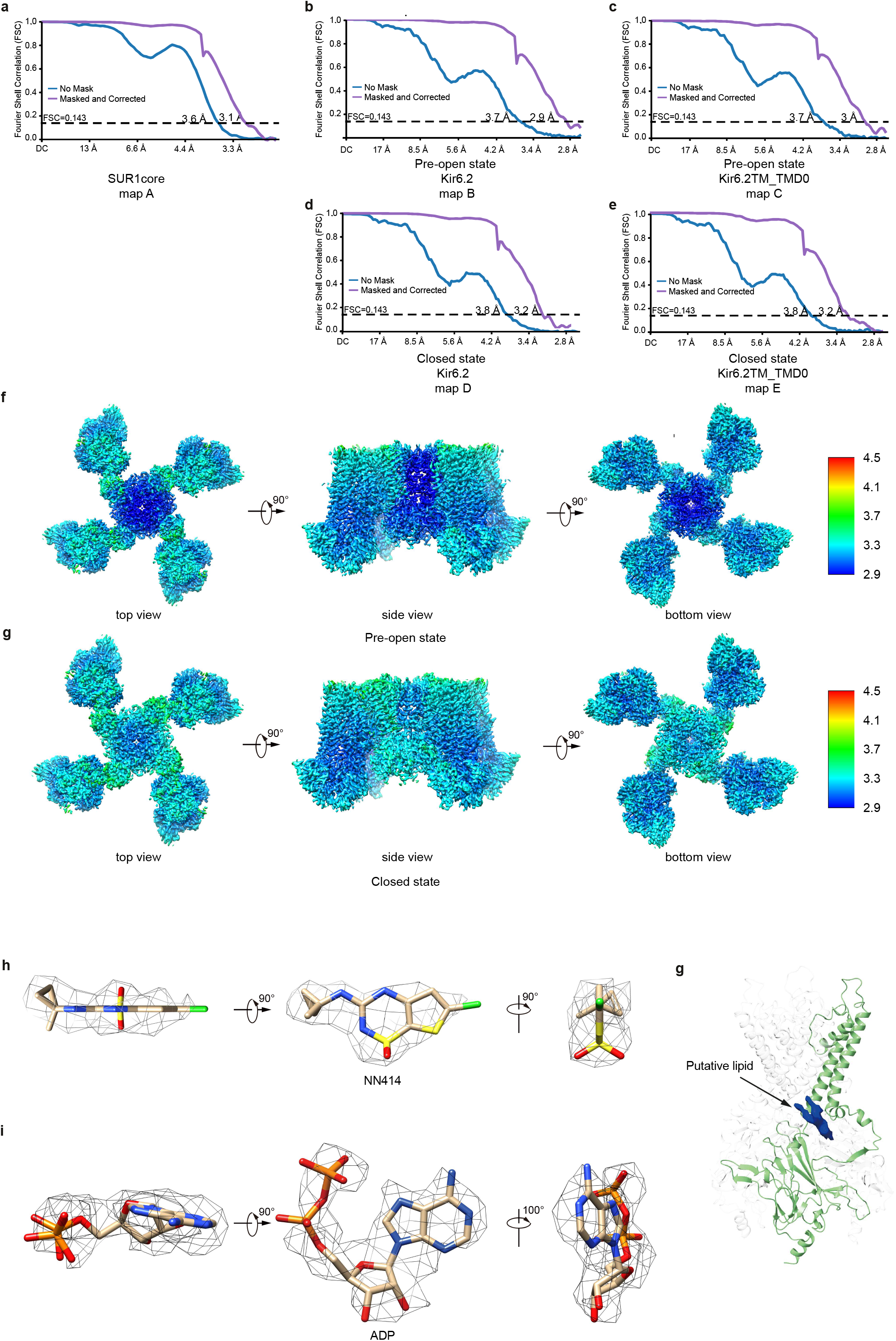
Electron densities of K_ATP_ H175K_cryoEM_. **a-e**, FSC curves of each focus refined map shown in Fig. S2. **f-g**, Local resolution estimation of the composite maps. **h**, Electron density maps of NN414 viewed from different angles. NN414 is shown as sticks. Electron density is shown as gray mesh. **i**, Electron density maps of ADP bound in Kir6.2 viewed from different angles. **j**, Electron density of putative lipid bound in Kir6.2 is shown in blue.

**Figure S4.**
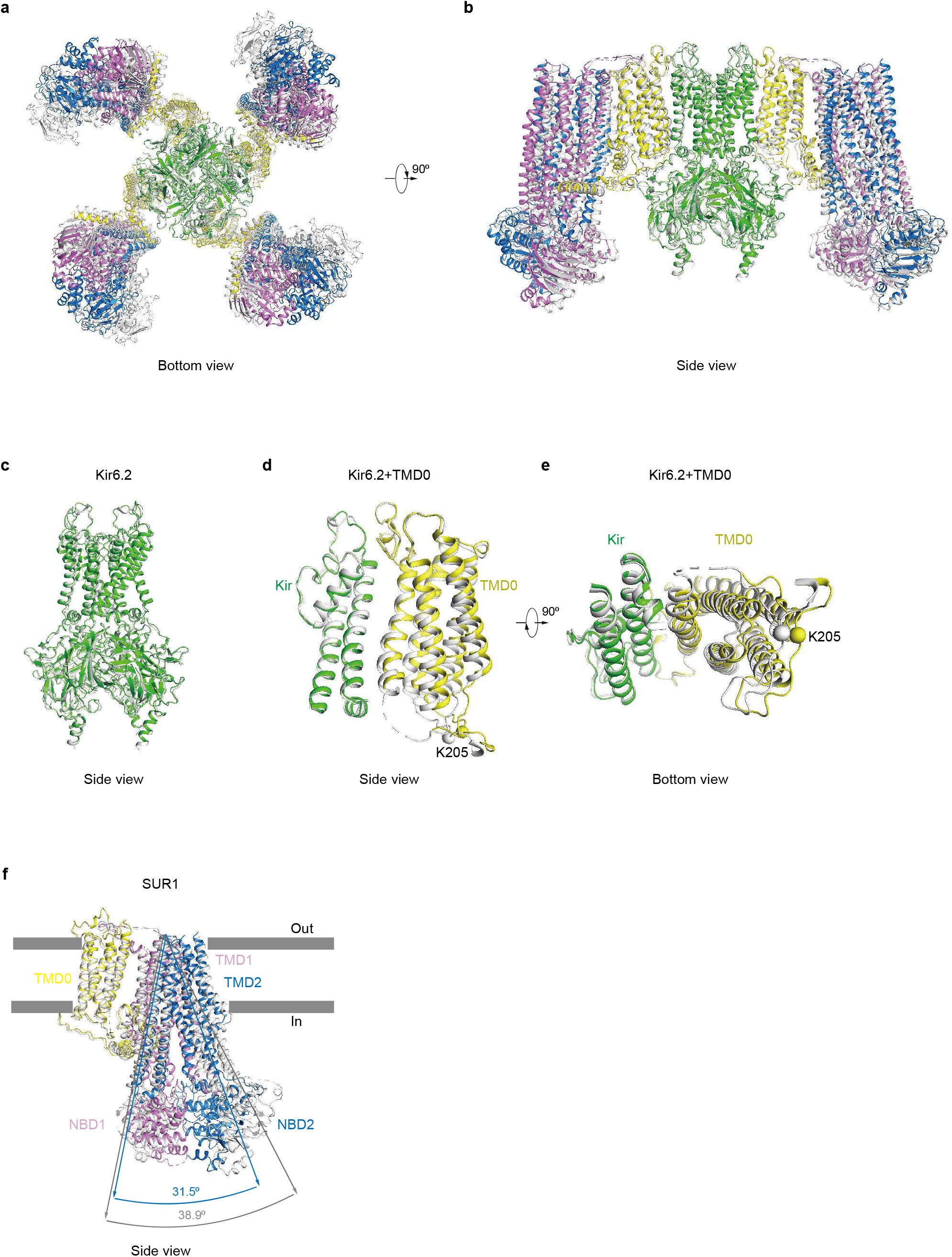
Structural comparison between the closed state of K_ATP_ H175K_cryoEM_ and K_ATP_ in the presence of ATP+RPG. **a-b,** Superposition of the closed state structure of H175K_cryoEM_ and K_ATP_ in the presence of ATP+RPG (PDB ID: 6JB1) by aligning the TMD of Kir6.2. The bottom view (**a**) and side view (**b**) are shown. H175K_cryoEM_ is colored according to Fig. 1. The structure of K_ATP_ in the ATP+RPG state is colored in gray. **c,** Structural comparisons of Kir6.2 in the closed state of H175K_cryoEM_ (green) and in the ATP+RPG state (gray). **d-e**, Structural comparisons of Kir6.2 and TMD0 in the closed state of H175K_cryoEM_ (colored) and in the ATP+RPG state (gray). The Cα positions of K205 are shown as spheres. **f,** Structural comparison of SUR1 in the closed state of H175K_cryoEM_ (colored) and in the ATP+RPG state (gray). Angles between M10 and M16 in the two structures are measured and labeled.

**Figure S5.**
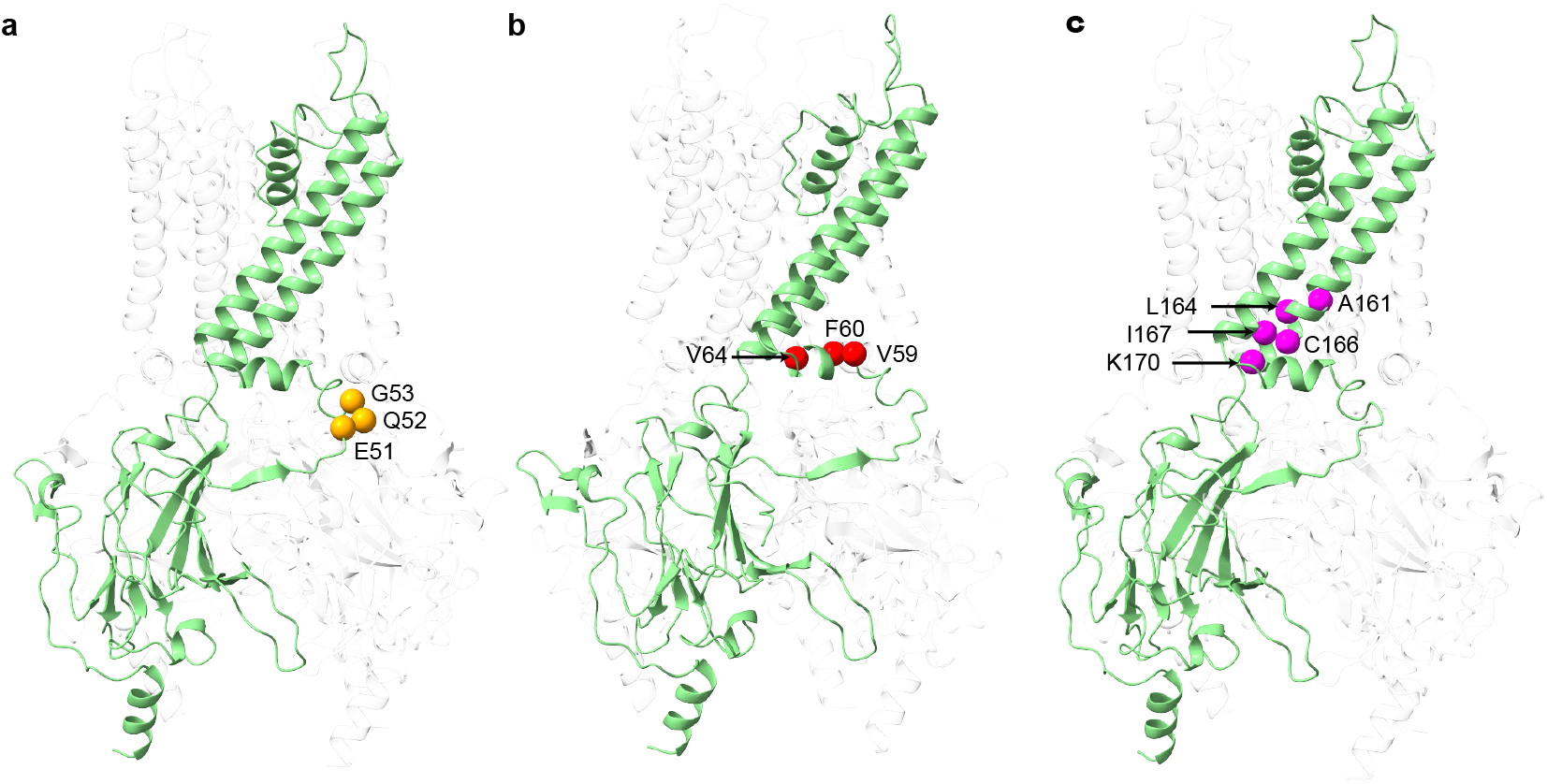
The spatial distribution of neonatal diabetes mutations mapped onto the structure of Kir6.2. Only mutations outside the ATP binding pocket of Kir6.2 are shown. **a,** Mutations on the βA-IH linker. Their Cα positions were shown as orange spheres. **b,** Mutations on IH helix (red spheres). **c**, Mutations on M2 helix (magenta spheres).

**Table S1.**
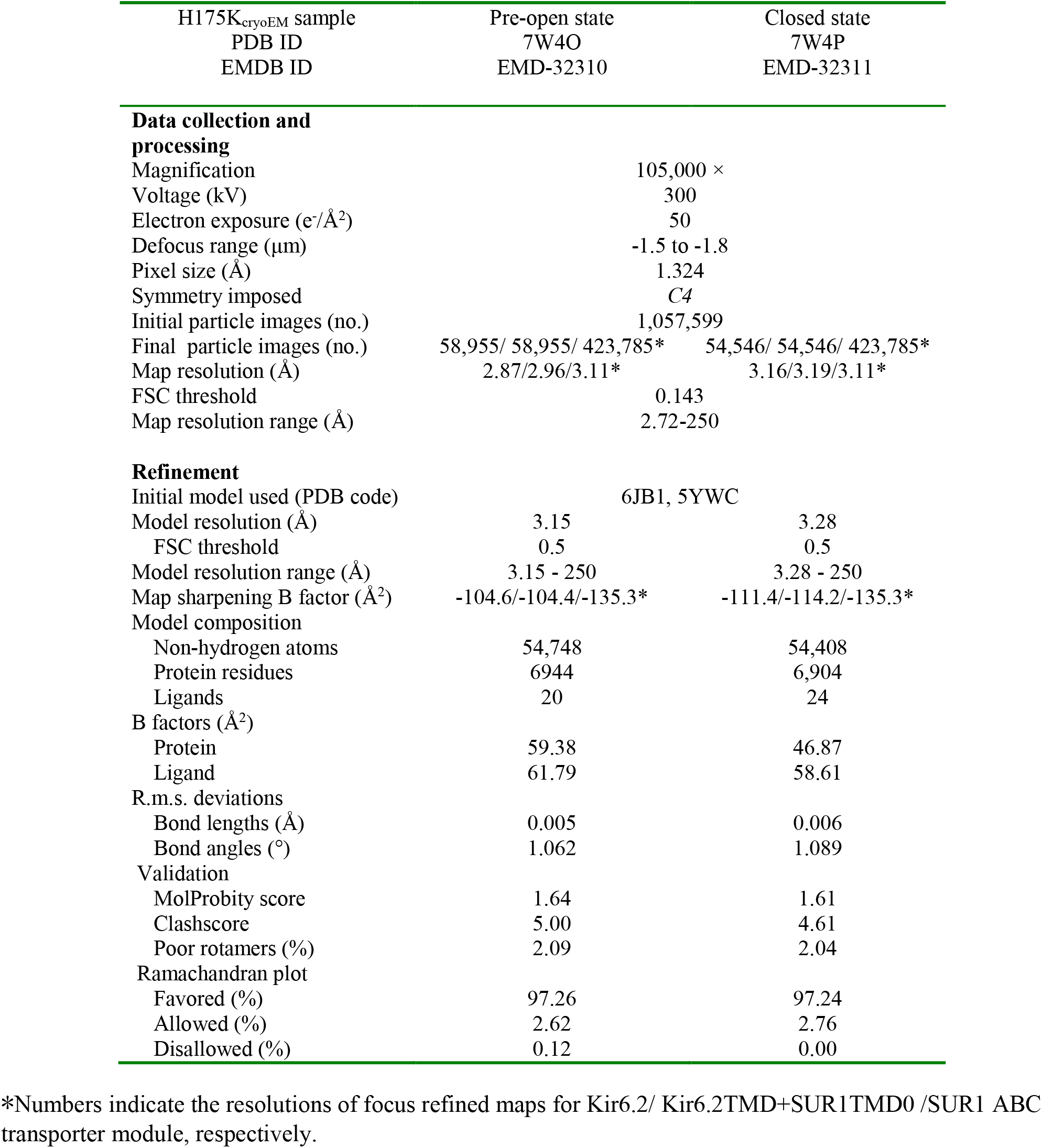
Cryo-EM data collection, refinement and validation statistics

